# Epidemiology and virological characteristics of the emerging SARS-CoV-2 variant PJ.2.1

**DOI:** 10.64898/2026.10.08.757354

**Authors:** Markus Hoffmann, Lu Zhang, Nianzhen Chen, Nicole Moor, Inga Nehlmeier, Luise Graichen, Anna Sophie Moldenhauer, Alexandra Dopfer-Jablonka, Christine Happle, Georg M N Behrens, Stefan Pöhlmann

## Abstract

SARS-CoV-2 continues to circulate globally at high levels and to evolve into new variants. Recently, a new saltation variant harbouring a highly mutated spike protein, PJ.2.1, emerged in North America. Here, we provide a brief epidemiological and virological characterization of PJ.2.1 and cocirculating variants. We report that PJ.2.1 is spreading in the USA, Canada, and Spain and mainly infects individuals aged 60 years and older, as do most other variants, with the exception of BA.3.2, which preferentially infects individuals aged 0-9 years. PJ.2.1 was less susceptible to inhibition by an antibody against ACE2 than were the other variants, indicating increased ACE2 binding. In contrast, PJ.2.1 did not show augmented evasion of antibodies from vaccinees who had received the LP.8.1-adapted vaccine compared with most other variants analysed. These results suggest that augmented ACE2 affinity rather than increased antibody evasion might drive the current, rapid spread of PJ.2.1.

## Introduction

SARS-CoV-2 continues to circulate globally and remains responsible for substantial morbidity and mortality. New COVID-19 waves are driven by variants arising through either antigenic drift or shift. Drift variants differ from their precursors by only a few mutations located mainly in the spike protein, which typically reduce sensitivity to antibody-mediated neutralization. Shift variants, also termed saltation variants, are highly mutated viruses thought to have evolved during persistent infections before re-entering global circulation. As a consequence of the large number of mutations in the spike protein, antibody neutralization may be markedly reduced. A novel saltation variant, PJ.2.1, was identified in May 2026 in Canada and is currently spreading. Here, we present a preliminary epidemiological and virological characterization of PJ.2.1 and cocirculating variants.

## Results and Discussion

PJ.2.1 originated from MC.10.1.7, a JN.1-descendant that circulated until May 2025, and was first detected in Ontario, Canada, on May 2, 2026 (Supplementary Figure 1). Subsequently, additional PJ.2.1 infections were reported from Ontario and New Brunswick. On June 30, 2026, PJ.2.1 was first detected in the USA in a patient in New York City followed by detections in California, Arizona, Maryland, New Jersey and Florida (Supplementary Figure 4 and 5). At present, based on wastewater analyses, PJ.2.1 is the fastest growing variant in the USA ^1^. Recently, PJ.2.1 has also been detected in Singapore (3 sequences), Australia (1 sequence), and Spain. In Spain, apparently local transmission chains were established and detections in Catalunya, Asturias, Murcia, Basque Country and Madrid were reported (Supplementary Figure 4 and 5).

**Figure 1:**
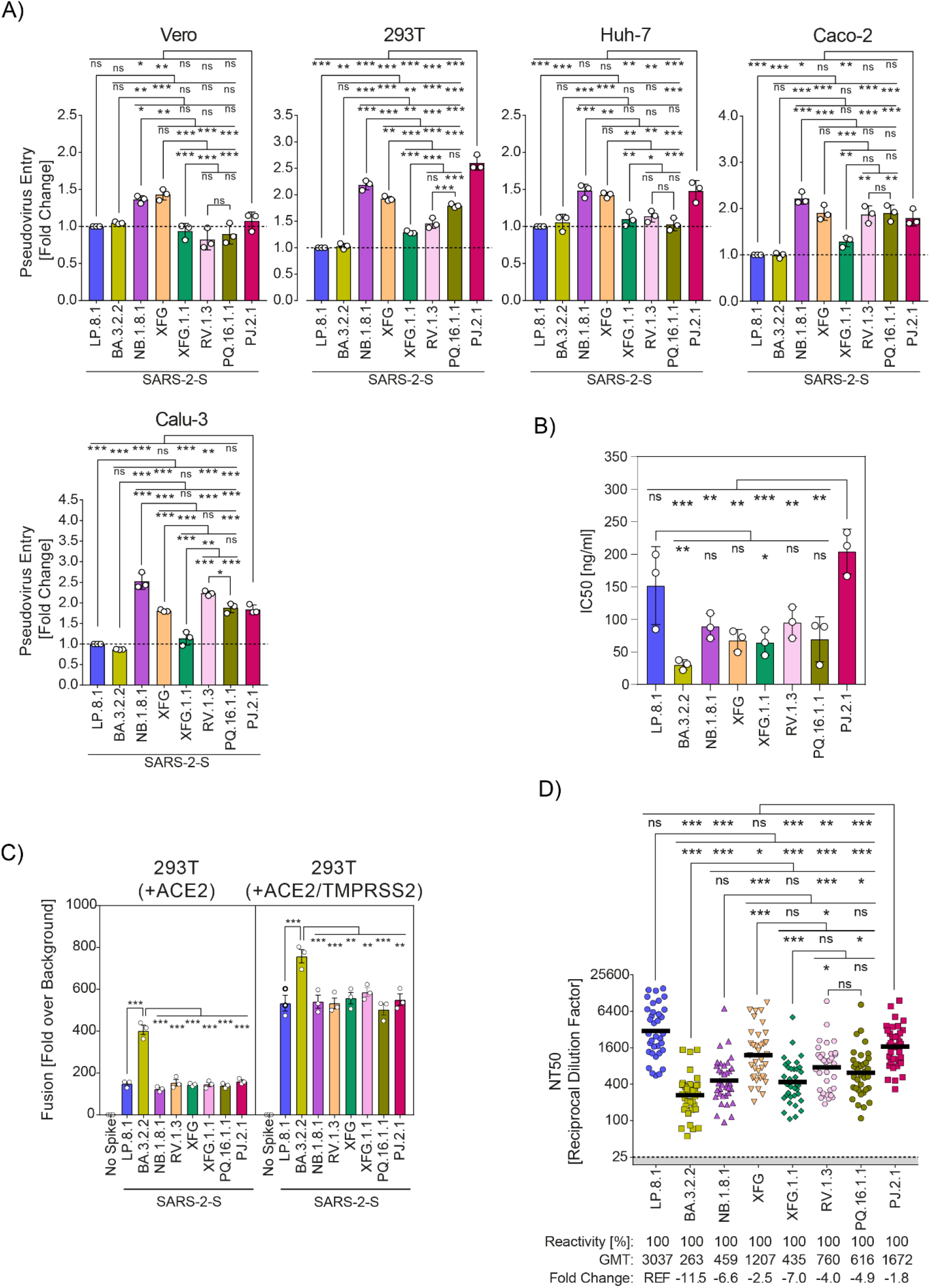
Host cell entry, ACE2 usage and cell-cell fusion driven by the PJ.2.1 S protein. **(A)** Vero, 293T, Huh-7, Caco-2, and Calu-3 cells were inoculated with pseudovirus particles containing the indicated S proteins. At 16-18 h postinoculation, entry efficiency was analysed by quantification of firefly luciferase in cell lysates and entry was normalised to particles bearing the SARS-CoV-2 LP.8.1 S protein (set as 1). Graphs show average (mean) data from three biological replicates, each conducted with four technical replicates. Error bars represent the standard error of the mean (SEM). Additional information can be found in supplementary figure 2A. **(B)** Vero cells were preincubated with different concentrations of ACE2-blocking antibody before inoculation with pseudovirus particles containing the indicated S proteins. At 16-18 h postinoculation, entry efficiency was analysed by quantification of firefly luciferase in cell lysates, normalised to entry in the absence of anti-ACE2 antibody (0% inhibition) and anti-ACE2 concentrations causing half-maximal inhibition (inhibitory concentration 50, IC50) were calculated based on a nonlinear regression model. The graph shows average (mean) IC50 values from three biological replicates, each conducted with four technical replicates. Error bars represent the standard deviation (SD). Additional information can be found in supplementary figure 2B. **(C)** Effector 293T cells expressing the indicated S proteins (or empty vector, control) and the β-galactosidase alpha fragment were mixed and co-cultivated with target 293T cells expressing human ACE2 and the β-galactosidase omega fragment (+ACE2), or target 293T cells expressing human ACE2, the β-galactosidase omega fragment and human TMPRSS2 (+ACE2/TMPRSS2). Following co-cultivation, S protein-driven cell-cell fusion was analysed by measuring the activity of reconstituted β-galactosidase activity and fusion was normalised to signals obtained for effector cells expressing no spike protein that were cultivated with either +ACE2 or +ACE2/TMPRSS2 target cells. The graph shows average (mean) data from three biological replicates, each conducted with four technical replicates. Error bars represent the SEM. Additional information can be found in supplementary figure 2C. **(D)** Pseudovirus particles containing the indicated S proteins were preincubated with culture medium containing serially diluted plasma from individuals vaccinated with Corminaty LP.8.1 (n = 36) before samples were inoculated onto Vero cells. At 16-18 h postinoculation, entry efficiency was analysed by quantification of firefly luciferase in cell lysates, normalised to particles incubated with culture medium without plasma (0% inhibition) and plasma dilution factors causing half-maximal inhibition (neutralising titre 50, NT50) were calculated based on a nonlinear regression model. The graph shows individual NT50 values (symbols) and geometric mean titres (GMT, black lines) from a single biological replicate conducted with four technical replicates. Numerical values below the graph indicate response rates (fraction of plasma samples showing neutralising activity), GMT values, mean fold change values in GMTs relative to pseudovirus particles containing the SARS-CoV-2 LP.8.1 S protein. Additional information can be found in supplementary table 1 supplementary figure 3. For panels A-C, statistical significance was assessed by one-way analysis of variance with Tuckey’s multiple comparison test while Friedman test with Dunn’s multiple comparison test was used for panel D (ns [not significant], p > 0.05; *, p ≤ 0.05; **, p ≤ 0.01; ***, p ≤ 0.001; ****, p ≤ 0.0001).

We also analysed variants BA.3.2.2, LP.8.1, NB.1.8.1 and XFG, which emerged in 2024/2025 and gave rise to the currently circulating variants PQ.16.1.1 (derived from NB.1.8.1), XFG.1.1 (derived from XFG), RV.1.3 (derived from XFG) (Supplementary Figure 1)^2-4^. BA.3.2.2 frequency was high in all areas analysed in May and June 2026 and was subsequently declining (Supplementary Figure 4). NB.1.8.1-derived variants dominated in Australia and Asia while XFG-derived variants dominated in Europe and North America. Substantial PQ.16.1.1 circulation was noted in Asia while RV.1.3 and XFG.1.1 spread was readily detectable in Europe and North America (Supplementary Figure 4). The analysis of age distribution of infected individuals revealed that BA.3.2.2 preferentially infected age group 0-9, as expected ^5^, while the remaining variants, including PJ.2.1, preferentially targeted individuals aged 70 or older (Supplementary Figure 4).

We next investigated whether PJ.2.1 differs from the other variants regarding cell line tropism, ACE2-dependence and sensitivity to antibody-mediated neutralization. All spike proteins examined mediated efficient fusion with cells transfected to express the ACE2 receptor and fusion activity was augmented by coexpression of the spike protein-activating cellular protease TMPRSS2 (Figure, panel C, and Supplementary Figure 2). Membrane fusion activity of BA.3.2.2 spike was highest, the activity of the remaining spike proteins was comparable. For the investigation of cell line tropism, we employed a well-established pseudotyping system ^6^ and cel lines, which are frequently used for SARS-CoV-2 research. We found that control particles pseudotyped with the glycoprotein of the vesicular stomatitis virus (VSV_pp_) entered all cell lines efficiently while particles without S protein (No Spike) did not, as expected (Figure, panel A, and Supplementary Figure 2). All spike proteins examined facilitated robust cell entry but some differences were noted. NB.1.8.1_pp_ and XFG_pp_ entered all cell lines with roughly 1.5-2.5-fold higher efficiency than LP.8.1_pp_, which served as reference, while BA.3.2.2_pp_ and XFG.1.1_pp_ entry was comparable to that of LP.8.1_pp_. Entry of RV.1.3_pp_ and PQ.16.1.1_pp_ into Caco-2 (colon) and Calu-3 (lung) cell lines was about 2-fold augmented relative to LP.8.1_pp_ (Figure, panel A, and Supplementary Figure 2). Finally, entry of PJ.2.1_pp_ into 4 out of the 5 cell lines tested was about 1.5-to 2.5-fold increased as compared to LP.8.1_pp_, and PJ.2.1_pp_ entered 293T (kidney) cells, which are known to express low levels of endogenous ACE2, with the highest efficiency of all pseudotyped particles studied.

We next employed an anti-ACE2 antibody to determine the efficiency of spike protein binding to ACE2. BA.3.2.2_pp_ were most efficiently inhibited, as evidenced by the lowest inhibitory concentration 50% measured (IC50) (25 ng/ml, Figure, panel B, and Supplementary Figure 2), as expected ^4^. NB.1.8.1pp, XFG_pp_, XFG.1.1_pp_, RV.1.3_pp_ and PQ.16.11_pp_ were comparably inhibited (IC50 between 60 and 100 ng/ml) and were more sensitive to inhibition than LP.8.1_pp_ (IC50 of 150 ng/ml). Finally, PJ.2.1 was least sensitive to inhibition by the anti-ACE2 antibody (IC50 of 200 ng/ml), indicating augmented ACE2 binding efficiency (Figure, panel B) and Supplementary Figure 2).

We finally tested neutralization by antibodies in donors present 2 weeks after receiving Comirnaty LP.8.1 that was recommended for use in the 2025/2026 COVID-19 season. LP.8.1_pp_ were sensitive to neutralization and served as reference (Figure, panel D, and Supplementary Table 1 and Supplementary Figure 2 and 3). Neutralization of BA.3.2.2_pp_ was least efficient and 11.5-fold reduced as compared to LP.8.1_pp_. NB.1.8.1_pp_ and XFG.1.1_pp_ also showed considerable antibody evasion with neutralization efficiency being 6.6- and 7-fold reduced, respectively, as compared to LP.8.1_pp_. Finally, neutralization of XFG_pp_, RV.1.3_pp_ and PQ16.11_pp_ was 2.5-, 4.0 and 4.9-fold reduced while neutralization of PJ.2.1_pp_ was only 1.8-fold lower than that of LP.8.1_pp_ (Figure, panel D, and Supplementary Figure 2 and 3).

Our results show that the global SARS-CoV-2 circulation is characterized by a heterogenous spectrum of variants with substantial regional differences. The emerging saltation variant PJ.2.1 does not exhibit high antibody evasion and in the light of the similar neutralization of XFG and PJ.2.1 one can speculate that the XFG-adapted vaccine approved for the 2026/2027 COVID-19 season might boost antibodies that efficiently neutralize PJ.2.1. In the light of the moderate antibody evasion of PJ.2.1, the rapid spread of this variant might be driven primarily by its increased ACE2 binding, which may enhance infection of cells expressing low levels of ACE2 and thereby augment transmissibility. Whether a potentially augmented spectrum of target cells also has an impact on disease presentation, including invasion of brain and gut, remains to be investigated.

