## Supplemental Information for "Epidemiology and virological characteristics of the emerging SARS-CoV-2 variant PJ.2.1"

### Methods

#### Cell culture

293T cells (human, female, kidney; ACC-635, DSMZ; RRID: CVCL 0063), Huh-7 (human, male, liver; JCRB0403, JRCB; RRID:CVCL_0336, kindly provided by Thomas Pietschmann) and Vero cells (African green monkey, female, kidney; CRL-1586, ATCC; RRID: CVCL 0574, kindly provided by Andrea Maisner) were maintained in Dulbecco's modified Eagle medium (DMEM; PAN-Biotech), which was supplemented with 10% (v/v) fetal bovine serum (FBS, Biochrom) and 1% (v/v) penicillin/streptomycin solution (pen/strep, PAN-Biotech). In addition, Calu-3 (human, male, lung; HTB-55, ATCC; RRID: CVCL_0609, kindly provided by Stephan Ludwig) and Caco-2 cells (human, male, colon; HTB-37, ATCC, RRID: CVCL_0025) were maintained in DMEM/F-12 medium with Nutrient Mix (Thermo Fisher Scientific) and minimum essential medium (Thermo Fisher Scientific), respectively, and media were supplemented with 10% (v/v) FBS, 1% (v/v) pen/strep solution, 1% (v/v) non-essential amino acid solution (PAA), and 1 mM sodium pyruvate (PAN-Biotech). Culturing of cell lines occurred at a temperature of 37 °C in a humidified atmosphere containing 5% $CO_2$. Cell line authentication was achieved by STR profiling, PCR amplification and sequencing of a fragment of the cytochrome c oxidase gene, and by analysis of cell morphology and growth characteristics of the cell lines under study. Further, the mycoplasma-free status of the cell lines was regularly confirmed by PCR. Transfection of 293T cells was achieved by calcium phosphate precipitation.

#### Expression plasmids and sequence analysis

The expression plasmids pCAGGS-DsRed, pCAGGS-VSV-G, pCG1-SARS-2-SΔ18 (LP.8.1) (EPI_ISL_19553000), pCG1-SARS-2-SΔ18 (BA.3.2.2) (EPI_ISL_19893762), pCG1-SARS-2-SΔ18 (NB.1.8.1) (EPI_ISL_19826033), pCG1-SARS-2-SΔ18 (XFG) (EPI_ISL_20053327), pQCXIBl-cMYC-hTMPRSS2, pQCXIP-β-galactosidase alpha fragment, and pQCXIP-β-galactosidase omega fragment have been described previously [1-7]. The pCG1 expression vector was generously provided by Roberto Cattaneo. Expression plasmids pCG1-SARS-2-SΔ18 (XFG.1.1) (EPI_ISL_20484922) and pCG1-SARS-2-SΔ18 (RV.1.3) (EPI_ISL_20485417) were generated by introducing the respective mutations (XFG.1.1: E96D, R445P, W452R; RV.1.3: L335F, A1174V, S446N) into plasmid pCG1-SARS-2-SΔ18 (XFG) through overlap-extension (primer sequences available upon request). Similarly, the expression plasmid pCG1-SARS-2-SΔ18 (PQ.16.1.1) (EPI_ISL_20489970) was generated by introducing the respective mutations (D253G, N417T, D420N, K478T) into plasmid pCG1-SARS-2-SΔ18 (NB.1.8.1). Finally, the expression plasmid pCG1-SARS-2-SΔ18 (PJ.2.1) (EPI_ISL_20503474) was generated by Gibson assembly. For this, linearized pCG1 plasmid (digested with BamHI and XbaI) was mixed with five synthetic DNA fragments sharing overlapping regions of 40-80 base pairs (Thermo Fisher Scientific) and GeneArt Gibson Assembly HiFi Master Mix (Thermo Fisher Scientific), and heated to 50 °C for 1 h. Next, the mixtures were transformed into One Shot™ OmniMAX™ 2 T1R chemically-competent *Escherichia coli* bacteria (Thermo Fisher Scientific). Screening of clones was done by colony PCR targeting the S protein encoding open reading frame. Of note, Information on SARS-CoV-2 lineages and associated mutations was retrieved from the GISAID database (Global Initiative on Sharing All Influenza Data; https://gisaid.org/) and the GitHub cov-lineages/pango-designation resource (https://github.com/cov-lineages/pango-designation/blob/master/lineage_notes.txt). All S protein sequences were codon-optimized (for human codon usage) and contained a truncation of the last 18 amino acid residues at the C-terminus. Further, sequence integrity of all constructs was confirmed by Sanger sequencing using a commercial service (Microsynth, SeqLab).

**Pseudovirus particle production and infection of target cells**

Pseudovirus particle preparation was done using a previously published protocol [5,8]. For this, 293T cells were transfected with expression plasmid encoding the respective S protein, VSV-G (positive control), or DsRed (negative control). Following an incubation period of 24 h, the cell culture supernatant was aspirated and cells were washed with PBS. Next, cells were inoculated with VSV-G-*trans*-complemented VSV*ΔG (FLuc) [9], a replication-restricted vesicular stomatitis virus (VSV) whose genome lacks the open reading frame for the VSV glycoprotein and instead codes for two independent open reading frames encoding eGFP and firefly luciferase (kindly provided by Gert Zimmer). The inoculum was aspirated after an incubation period of 1 h at 37

°C before cells were washed with PBS and further incubated with medium containing anti-VSV-G antibody (supernatant from I1-hybridoma cells; ATCC no. CRL-2700). Of note, cells initially transfected with expression plasmid for VSV-G were incubated with antibody-free medium. Pseudovirus-containing supernatants were harvested at 16-18 h postinoculation. For this, supernatants were cleared from cellular debris by centrifugation (4,000 x g, 10 min) and clarified supernatants were further aliquoted and stored at -80 °C.

To investigate the cell line tropism and entry efficiency of S protein-bearing pseudovirus particles, target cells grown in 96-well plates were inoculated with equal volumes of pseudovirus particles and incubated for 16-18 h. For analysis of the impact of anti-ACE2 antibody-mediated blockade of S protein-driven cell entry, Vero cells were preincubated for 30 min at room temperature with different concentrations of ACE2-blocking antibody (Sino Biologicals, 10108-MM36) diluted in culture medium (concentrations: 5,000, 1,000, 200, 40, 8, 1.6 ng/ml) or culture medium without antibody as control (0% inhibition). Next, pseudovirus particles were added and cells were further incubated. The efficiency of pseudovirus entry was quantified at 16-18 h post inoculation by quantification of the activity of virus-encoded firefly luciferase (as described below) and anti-ACE2 blockade efficiency was determined based on the relative inhibition of entry compared to samples without antibody (set to 0% inhibition).

Entry efficiency of pseudovirus particles was analysed through quantification of the activity of virus-encoded firefly luciferase in cell lysates. For this, culture medium was aspirated and cells were incubated for 30 minutes at room temperature with PBS containing 0.5% Tergitol 15-S-9 (Carl Roth) to achieve cell lysis. The cell lysates were then transferred to white 96-well plates. Finally, luciferase substrate (Beetle-Juice, PJK) was added and luminescence was recorded with a Hidex Sense plate luminometer (Hidex).

**Cell-cell fusion assay**

S protein-mediated cell-cell fusion was analysed according to a previously published protocol [10]. For this, effector 293T cells were co-transfected with expression plasmids for the respective S protein (or empty vector) and the β-galactosidase alpha fragment, while target 293T cells were co-transfected with expression plasmids for human ACE2, the β-galactosidase omega fragment, and either empty plasmid (-TMPRSS2 condition) or expression plasmid for human TMPRSS2 (+TMPRSS2 condition). Following an incubation period of 18 h, effector and target cells were washed with PBS and cultured in fresh medium for 6 additional hours. Next, target cells were washed with PBS and resuspended in fresh medium before being mixed with effector cells. This step was followed by co-cultivation of effector and target cells for 18 h to allow for S protein-driven cell-cell fusion, which was quantified following incubation of β-galactosidase substrate (Gal-Screen, Thermo Fisher Scientific) for 90 min and measurement of luminescence using a Hidex Sense plate luminometer (Hidex).

**Plasma samples and neutralisation assay**

We analysed plasma of 36 individuals that had been vaccinated with Corminaty LP.8.1 (30 µg; BioNTech Manufacturing GmbH, Mainz, Germany) in September 2025 as part of the COVID-19 Contact (CoCo) Study (German Clinical Trial Registry, DRKS00021152), an ongoing, prospective observational study designed to monitor immunoglobulin G (IgG) levels and immune responses in healthcare professionals and individuals with potential SARS-CoV-2 exposure at Hannover Medical School [11]. Blood samples were collected on day 14 after vaccination, when robust antibody responses are detectable [12]. All plasma samples were pre-screened for the absence of an increase in antibodies against the SARS-CoV-2 nucleocapsid protein (anti-NCP) to exclude samples from individuals with a recent SARS-CoV-2 infection. None of the plasma samples had anti-NCP IgG levels indicative for a recent SARS-CoV-2 infection. Demographics (sex and age) as well as infection and vaccination history are summarised in **Table S1**. Before their use in neutralisation assays, the plasma samples were heat-inactivated at 56 °C for 30 min. Neutralisation assays were conducted according to a previously published protocol [5,8]. For this, pseudovirus particles were mixed with serially diluted plasma (diluted in culture medium: 1:25, 1:100, 1:400, 1:1,600, 1:6,400) and incubated for 30 min at 37°C. In addition, pseudovirus particles incubated with culture medium without plasma were included as controls. Following incubation, the mixtures were added to confluent monolayers of Vero cells that were grown in 96-well plates. The efficiency of pseudovirus entry was quantified at 16-18 h post inoculation by quantification of the activity of virus-encoded firefly luciferase (as described above) and neutralisation efficiency was determined based on the relative inhibition of entry compared to samples without plasma (set to 0% inhibition).

**Quantification and statistical analysis**

Data were analysed with Microsoft Excel (part of Microsoft Office LTSC Standard 2024, version 2408, Microsoft Corporation) and GraphPad Prism (version 10.4.2, GraphPad Software). Half-maximal inhibitory anti-ACE2 concentrations (inhibitory concentration 50, IC50) and plasma dilutions (neutralising titre 50, NT50) were calculated based on a nonlinear regression model. Assessment of statistical significance was performed by one-way analysis of variance (ANOVA) with Tuckey's multiple comparison test or Friedman test with Dunn's multiple comparison test. Only p values of 0.05 or lower were deemed as statistically significant (ns [not significant], $p > 0.05$; *, $p \leq 0.05$; **, $p \leq 0.01$; ***, $p \leq 0.001$; ****, $p \leq 0.0001$).

**Ethics statement**

This research was conducted as part of the COVID-19 Contact (CoCo) Study (DRKS00021152), a prospective observational study designed to monitor IgG levels and immune responses in healthcare professionals and individuals with potential SARS-CoV-2 exposure at Hannover Medical School. Approval for the collection and analysis of plasma samples was obtained from the Institutional Review Board of Hannover Medical School (approval no. 8973_BO_K_2020). The participants provided written informed consent and did not receive financial compensation for being involved in the study.

**Limitations of the study**

This study has limitations. First, we used S protein-bearing pseudovirus particles and immortalised cell lines instead of authentic SARS-CoV-2 isolates and primary cell cultures to study host cell entry, cell-cell fusion and neutralisation sensitivity. Second, the number of plasma samples is limited and thus does not allow for the analysis the impact of biological variables (e.g., age and sex) on neutralisation efficiency. Third, neutralising activity in plasma samples was analysed at two weeks after Corminaty LP.8.1 vaccination and neutralising antibody responses against specific SARS-CoV-2 lineages may evolve differentially over time. Lastly, given the diverse immunisation background (vaccination and infection) in the global population, our study provides only a small excerpt of lineage-specific neutralising activity and neutralising antibody responses may for example differ with respect to cohorts vaccinated with non-mRNA vaccines and/or cohorts from geographic regions with high circulation of distinct SARS-CoV-2 lineages.

**Role of the funding sources**

IN, SP, and MH conducted contract research (testing of vaccine serums for neutralising activity against SARS-CoV-2) for Valneva, which is unrelated to this work. GMNB was an adviser for Moderna, and SP was an adviser for BioNTech, which is unrelated to this work. AD-J was an adviser for Pfizer, which is unrelated to this work. All other authors declare no competing interests.

AD-J acknowledges funding by the European Social Fund (ZAM5-87006761), and together with GMNB, by the COVID-19-Research Network Lower Saxony (COFONI) through funding from the Ministry of Science and Culture of Lower Saxony in Germany (14-76103-184, project 4LZF23). SP acknowledges funding by the DZIF, associated partner site Göttingen. NC acknowledges funding by the Chinese Scholarship Council (202308310035). The funding sources had no role in the design and execution of the study, writing of the manuscript, or the decision to submit the manuscript for publication. The authors did not receive payment from any pharmaceutical company or other agency to write the manuscript. The authors were not precluded from accessing data in the study, and they accept responsibility for submitting it for publication.

**Author contributions**

Conceptualisation: MH, GMNB, SP; Formal analysis: MH, SP; Funding acquisition: NC, GMNB, SP; Investigation: MH, LZ, NC, NM, IN, LG, ASM; Resources: AD-J, CH, GMNB; Writing–original draft: MH, SP; Writing–review and editing: all authors. MH and SP have directly accessed and verified the underlying data reported in the manuscript. ChatGPT (OpenAI) was used to assist with grammar and style corrections, to generate a schematic illustration (Supplementary Figure

5) showing the geographic locations in North America and Spain where PJ.2.1 infections were detected, and to assist in designing Excel spreadsheets for epidemiological analyses. The authors independently verified the functionality and accuracy of the Excel spreadsheets and the analyses performed with them, verified the information depicted in Supplementary Figure 5 against the underlying data, and carefully reviewed and revised all AI-assisted text. The authors take full responsibility for the content of the manuscript.

#### Acknowledgements

We acknowledge the originating laboratories responsible for sample collection and the submitting laboratories that generated and shared genome data via GISAID, which formed the basis of this study. We are further thankful to Roberto Cattaneo, Stephan Ludwig, Andrea Maisner, Thomas Pietschmann, and Gert Zimmer for providing reagents. Finally, we thank the participants of the CoCo Study for their contribution, as well as the entire CoCo Study team for their continued support. We thank the SARS-CoV-2 variant communities for sharing information and discussion on BlueSky and X (formerly Twitter). We particularly thank Ryan Hisner (@LongDesertTrain) for his insights into SARS-CoV-2 evolution.

### Supplementary Table 1

#### Demographics, and infection and vaccination history of plasma donors

| Variable | Vaccinees |
|---|---|
| Number of vaccinees | 36 |
| Age, Median (IQR) | 57 (16) |
| Sex, male (%) | 16 (44) |
| Prior COVID-19 vaccination (%) | 36 (100) |
| Prior JN.1 specific COVID-19 vaccination[1] (%) | 28 (78) |
| Number of COVID-19 vaccinations[2]: Median (IQR), range | 6 (1), 3-9 |
| Months since last COVID-19 vaccination[3]: Median (IQR), range | 13 (0), 10-45 |
| Prior SARS-CoV-2 infection, n (%) | 31 (86) |
| Number of SARS-CoV-2 infections: Median (IQR), range | 1 (1), 0-3 |
| Months since last SARS-CoV-2 infection[3]: Median (IQR), range | 35 (23), 13-59 |

[1]Missing information for three persons. [2]Missing information for two persons. [3]Missing information for four persons

#### Supplementary Figure 1

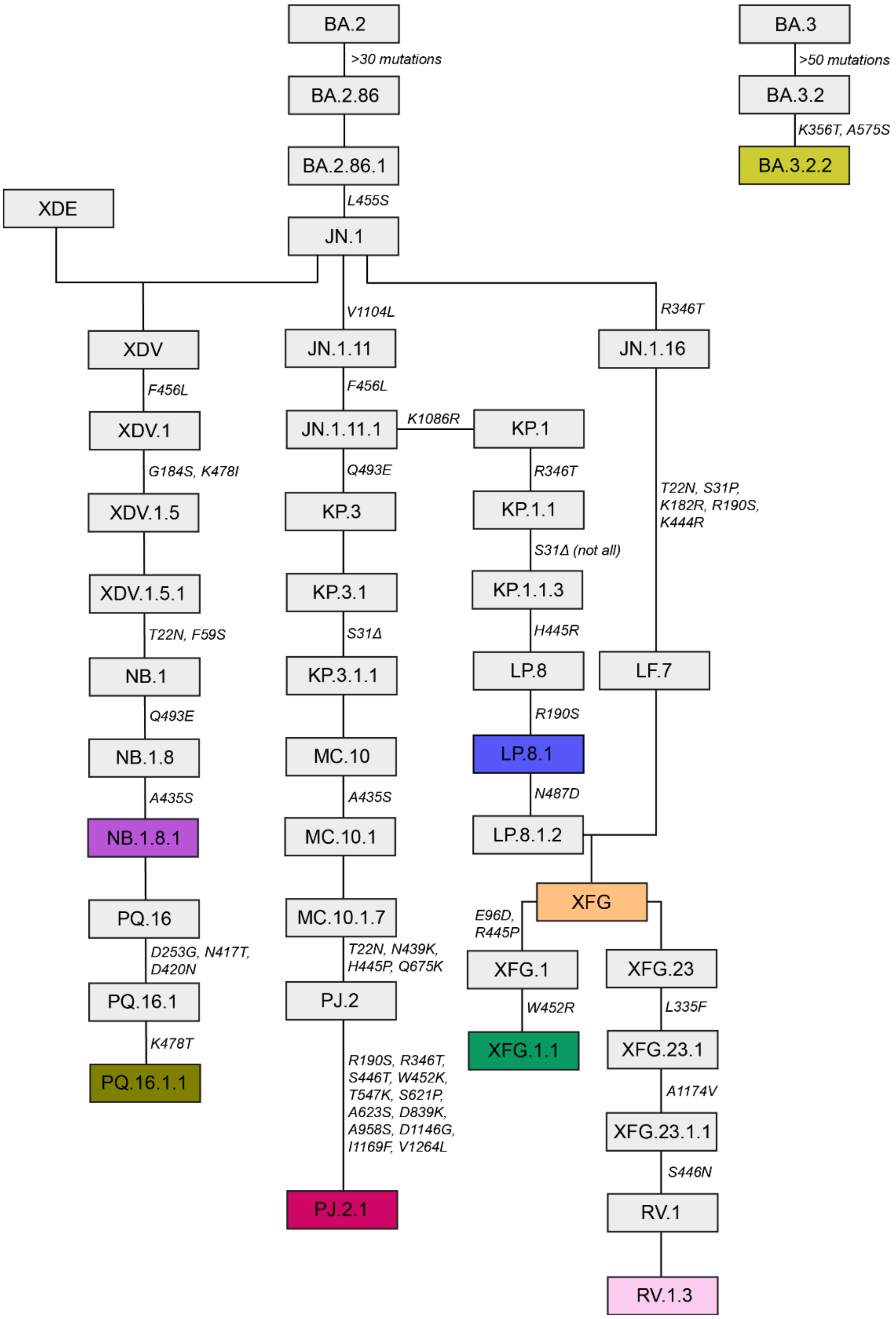

**Supplementary Figure 1: Lineage-specific S protein mutations.**

Acquisition of S protein mutations during SARS-CoV-2 lineage evolution. The lineages under study are highlighted. Information on SARS-CoV-2 lineages and associated mutations was retrieved from the GISAID database and the GitHub cov-lineages/pango-designation resource.

### Supplementary Figure 2

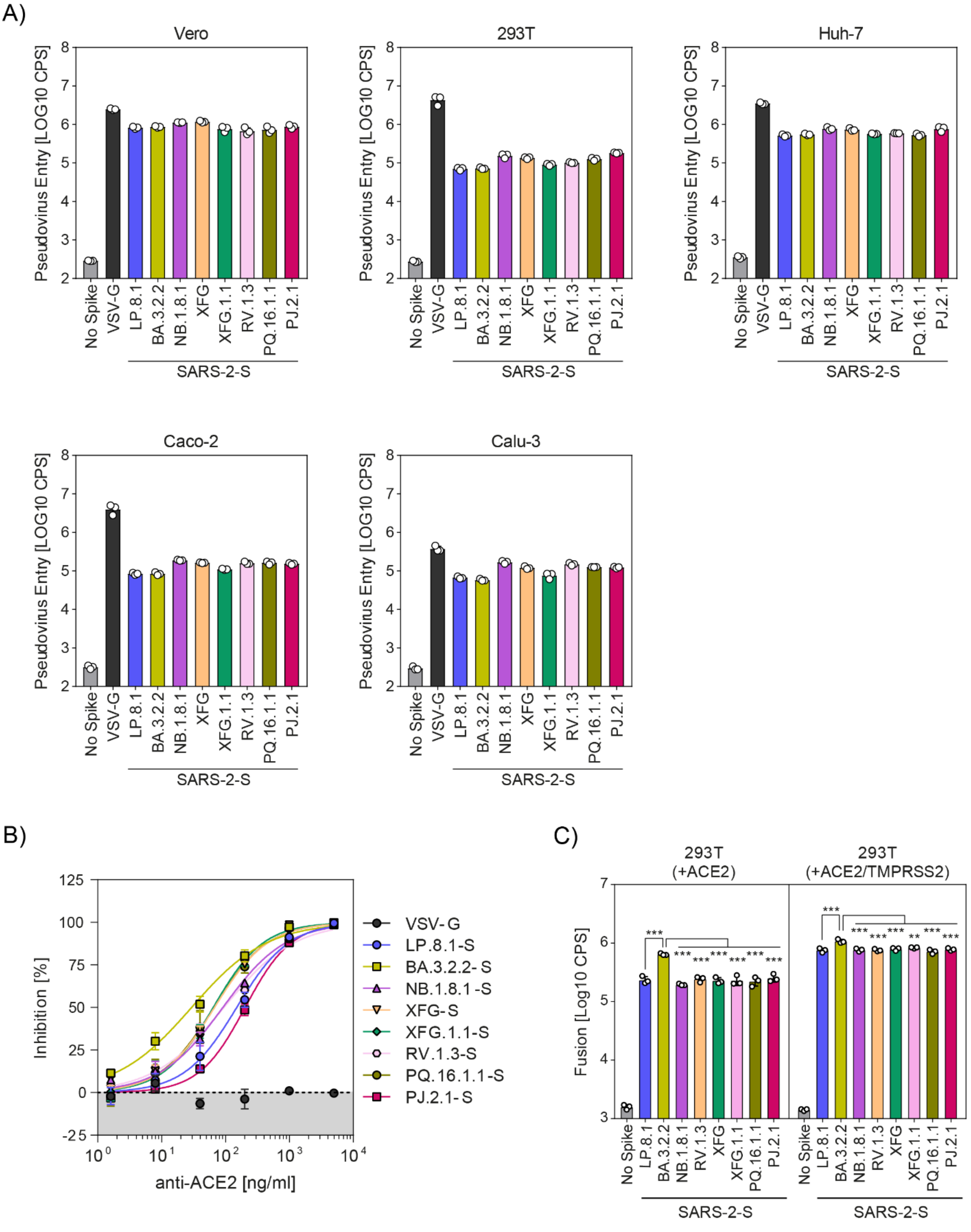

**Supplementary Figure 2: Host cell entry, ACE2 usage and cell-cell fusion driven by the S proteins under study.**

**(A)** Vero, 293T, Huh-7, Caco-2, and Calu-3 cells were inoculated with pseudovirus particles containing the indicated S proteins, VSV-G (positive control) or no viral glycoprotein (No Spike, negative control). At

16-18 h postinoculation, entry efficiency was analysed by quantification of firefly luciferase in cell lysates. Graphs show log10-transformed average (mean) data from three biological replicates, each conducted with four technical replicates. Error bars represent the standard deviation (SD). **(B)** Vero cells were preincubated with different concentrations of ACE2-blocking antibody before inoculation with pseudovirus particles containing the indicated S proteins or VSV-G (negative control). At 16-18 h postinoculation, entry efficiency was analysed by quantification of firefly luciferase in cell lysates. The graph shows average (mean) data from three biological replicates, each conducted with four technical replicates, for which entry efficiency was normalised using cells incubated in the absence of anti-ACE2 antibody as reference (0% inhibition). Error bars represent the standard error of the mean (SEM). **(C)** Effector 293T cells expressing the indicated S proteins (or empty vector, control) and the β-galactosidase alpha fragment were mixed and co-cultivated with target 293T cells expressing human ACE2 and the β-galactosidase omega fragment (+ACE2), or target 293T cells expressing human ACE2, the β-galactosidase omega fragment and human TMPRSS2 (+ACE2/TMPRSS2). Following co-cultivation, S protein-driven cell-cell fusion was analysed by measuring the activity of reconstituted β-galactosidase activity. The graph shows log10-transformed average (mean) data from three biological replicates, each conducted with four technical replicates. Error bars represent the SD. Statistical significance was assessed by one-way analysis of variance with Tuckey's multiple comparison test (ns [not significant], $p > 0.05$; *, $p \leq 0.05$; **, $p \leq 0.01$; ***, $p \leq 0.001$; ****, $p \leq 0.0001$).

### Supplementary Figure 3

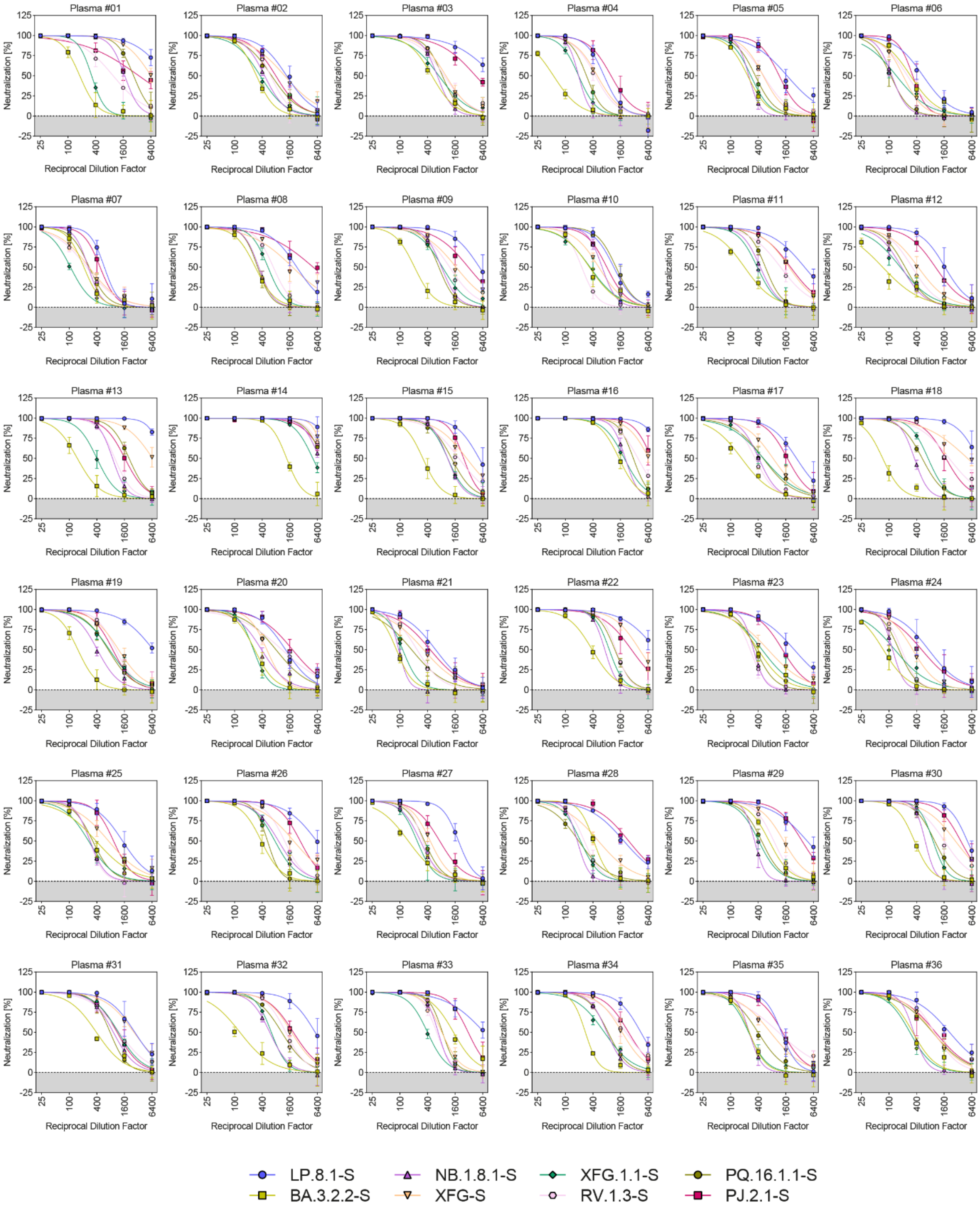

**Supplementary Figure 3: Individual neutralisation data.**

Plasma-specific neutralisation data for LP.8.1$_{pp}$, BA.3.2.2$_{pp}$, NB.1.8.1$_{pp}$, XFG$_{pp}$, XFG.1.1$_{pp}$, RV.1.3$_{pp}$, PQ.16.1.1$_{pp}$, and PJ.2.1$_{pp}$. Pseudovirus particles containing the indicated SARS-CoV-2 lineage S proteins were preincubated with serially diluted plasma samples obtained 14 days after vaccination with the mRNA vaccine Corminaty LP.8.1 and subsequently inoculated on Vero cells. Cell entry efficiency was analysed at 16-18 h postinoculation and normalised to control samples (pseudovirus particles incubated with culture medium without plasma, 0% inhibition). Graphs show average (mean) data from a single biological replicate conducted with four technical replicates and curves were fitted using a nonlinear regression model. Error bars represent the SEM.

### Supplementary Figure 4

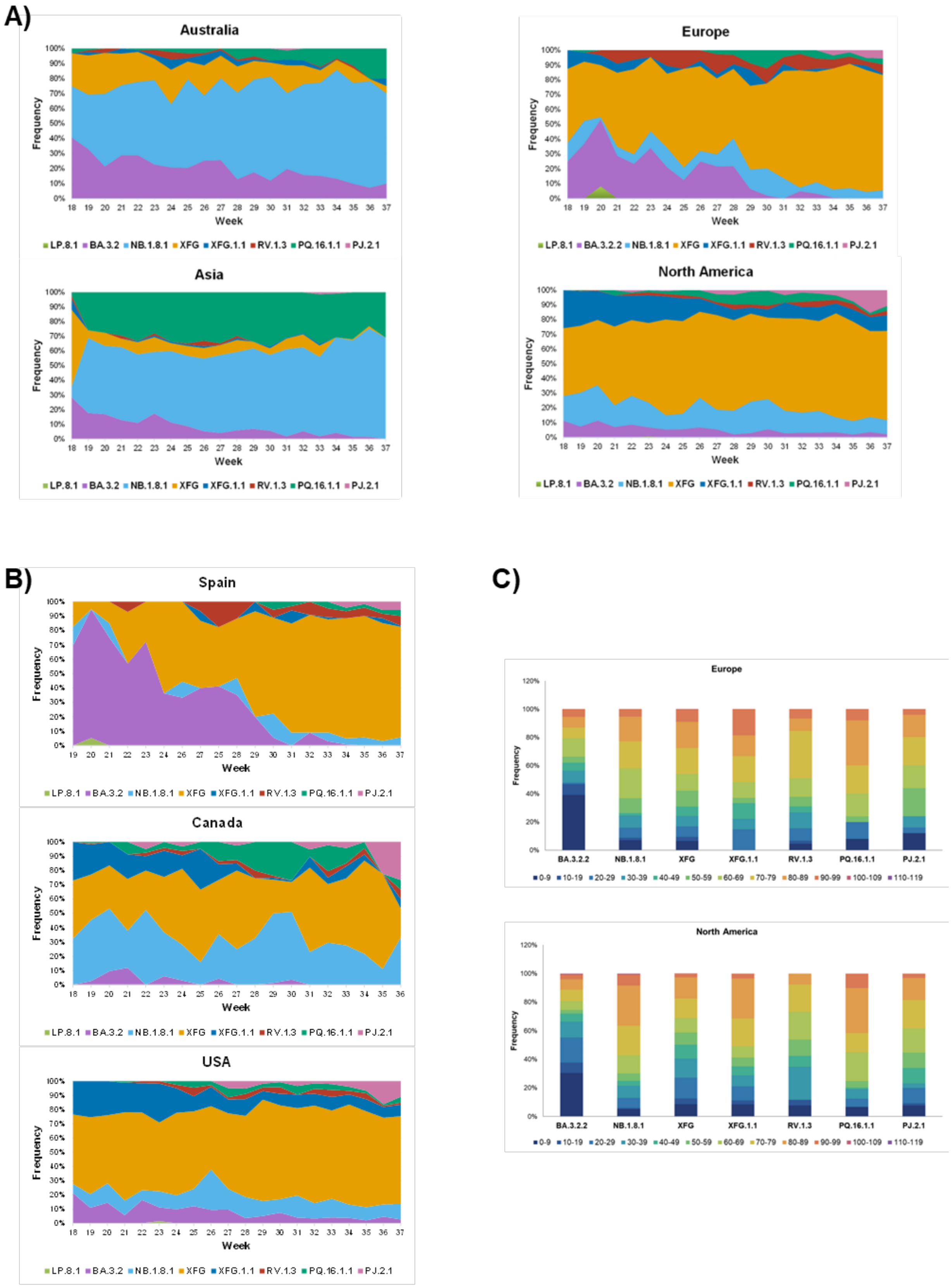

**Supplementary Figure 4: Frequency and age distribution of PJ.2.1 and co-circulating SARS-CoV-2 lineages.**

**(A)** SARS-CoV-2 sequences collected between May 1 and September 28 in the indicated geographical regions and deposited in GISAID were downloaded and classified by lineage. PJ.2.1 sequences were identified using Nextstrain. For sequences from Israel, only the month of sampling was available, with no specific sampling day provided. For the purposes of this analysis, the sampling date was therefore assumed to be the first day of the reported month. For each week and geographical region, the relative frequency of each indicated variant was calculated as a proportion of all available sequences collected during that week. **(B)** The analysis was conducted as described for panel (A) but sequences collected in the indicated countries were analysed. **(C)** SARS-CoV-2 sequences collected between May 1 and September 28 in Europe and North America and deposited in GISAID were downloaded, sorted for availability of donor age and age distribution per variant determined. The number of sequences analysed is shown above each bar. It should be noted that XFG.1.1 and RV.1.3 are subvariants of XFG, whereas PQ.16.1.1 is a subvariant of NB.1.8.1. Consequently, XFG.1.1 and RV.1.3 are included both individually and within the XFG-derived variant category and are therefore counted twice in the corresponding figure panel. The same applies to PQ.16.1.1 and NB.1.8.1

#### Supplementary Figure 5

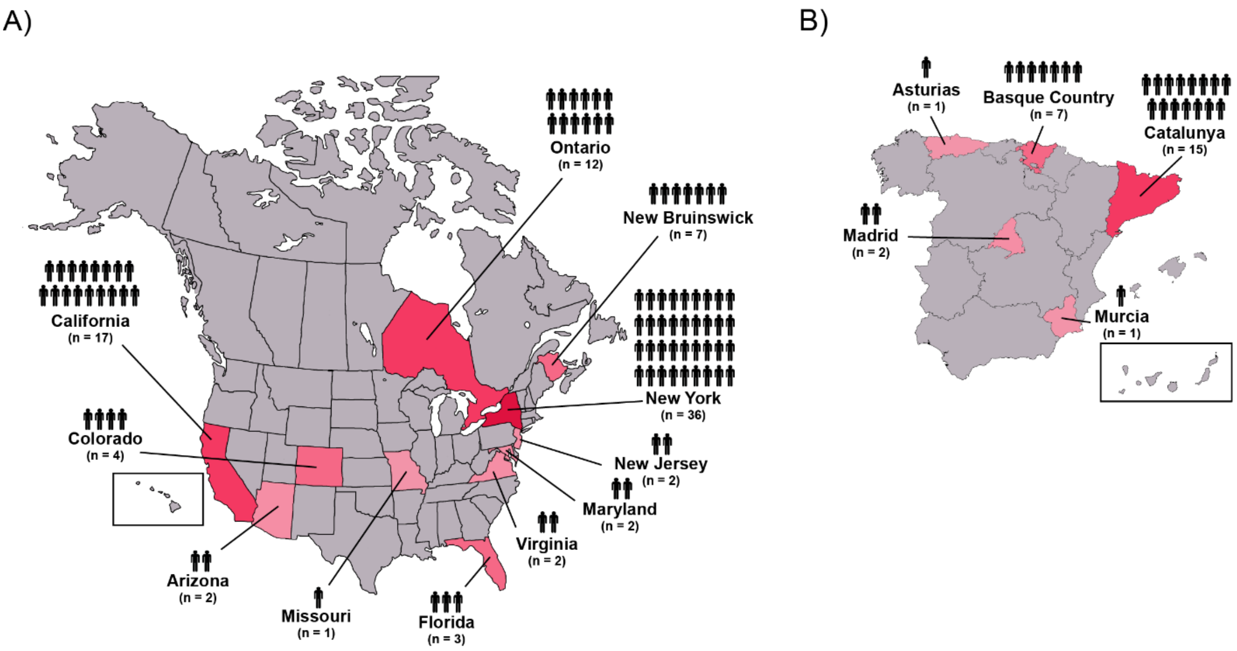

**Supplementary Figure 5: PJ.2.1 detections in North America and Spain.**

SARS-CoV-2 sequences collected between May 1 and September 28 in North America (upper panel) and Spain (lower panel) and deposited in GISAID were downloaded and PJ.2.1 sequences identified using the Nextstrain resource. The number and location of PJ.2.1 detections is indicated.
